# Island population demography: Breeding dynamics and drivers of Gotland’s iconic Golden Eagles

**DOI:** 10.1101/2024.11.02.621680

**Authors:** Navinder J Singh, Robin Olofsson, Aemilius Johannes van der Meiden, Andres Lopez-Painado, Johan Månsson

## Abstract

1. Raptor populations on islands are limited by resource availability and the dispersal possibilities for young birds, which are often determined by the size of the island. This leads to differences in population dynamics and viability compared to mainland populations. Human land use modifications on islands—such as agriculture, forestry, excessive hunting, and urban infrastructure development—may affect resource availability and increase risks to these populations, ultimately threatening their survival. Consequently, many island raptor populations have been dramatically reduced or driven to extinction and have never fully recovered. The conditions necessary for their long-term persistence remain uncertain.
2. Gotland, a large, human-dominated island located in the Baltic Sea, is home to one of the densest populations of Golden Eagles (*Aquila chrysaetos*) in the world. However, the drivers of population dynamics remain unknown, and many speculations exist that require empirical testing.
3. Approximately 86 Golden Eagle territories were identified and surveyed across Gotland, an island spanning approximately 3,200 km² (152 km long, 52 km wide, with an 800 km coastline). We investigated the spatial drivers of breeding dynamics in this eagle population, evaluating the effects of territorial habitat composition, overlap with White-tailed Eagles, prey density, and neighborhood effects on territorial productivity.
4. The average productivity was 0.41 fledglings per pair, which varied annually, with approximately 72% of territories occupied and 32% being successful. Despite significant variation in habitat composition across territories, spatial differences in productivity were primarily influenced by the proportion of coniferous forest (nesting habitat), access to coastal areas (greater prey diversity), the density of the main prey species (roe deer, *Capreolus capreolus*), and the reproductive status of neighboring territories in a year.
5. Several novel findings emerged: the role of roe deer as a potential prey species had been previously underappreciated, proximity to the coast was associated with increased productivity, and the variation in spatio-temporal reproductive dynamics across neighboring territories appears to influence overall population dynamics. This relationship warrants further study. We discuss the implications of these findings for the long-term conservation and persistence of this iconic island population and similar populations worldwide.

## Introduction

Raptors play a crucial role in ecosystems as top predators. Their presence and population dynamics on islands present unique ecological scenarios due to the isolation and limited resources, characteristic of insular environments (MacArthur & Wilson 1967). Hence, raptor populations on islands can be influenced by habitat availability, prey abundance, and inter as well as intraspecific competition differently as compared to mainland populations (White *et al*. 1971; Ferrer *et al*. 2011). Small islands may support only a few individuals due to restricted space and resources, leading to heightened competition and potential population fluctuations. Conversely, larger islands with diverse habitats may sustain larger and more stable raptor populations (Thibault *et al*. 1992; Donázar *et al*. 2005). Additionally, island populations may also face unique challenges such as genetic isolation and genetic drift, which can impact their long-term viability (Agudo *et al*. 2011; Sanz-Aguilar *et al*. 2015). These populations may undergo cyclical fluctuations in abundance, driven by factors such as prey availability, environmental conditions, and interspecific interactions. These population cycles can result in periods of population growth followed by crashes or declines (Barbraud *et al*. 2023).

The breeding ecology of raptors encompasses various aspects of their reproductive biology, nesting behavior, and parental care. Raptors select nesting sites on cliffs, trees, shrubs and human-made structures such as buildings or transmission towers depending on their availability (Bedrosian *et al*. 2020; Balotari-Chiebao *et al*. 2021). This may entail that raptors on islands may select a different range of nesting sites, according to the availability of habitats (Evans *et al*. 2010; Di Vittori & López-López 2014). The availability and suitability of nesting sites influences the breeding success and population dynamics, whereas the timing of breeding can be influenced by factors such as prey availability, temperature, precipitation, and photoperiod (Katzner *et al*. 2012; Miller *et al*. 2017; Maynard *et al*. 2024).

Therefore, breeding phenology on islands may differ from those on the mainland due to local environmental conditions and resource availability (Maynard *et al*. 2022; Gómez-López *et al*. 2023). In terms of resources, island ecosystems typically have fewer prey species compared to mainland areas (Amar *et al*. 2003; Balza *et al*. 2020). As a result, island predators may specialize on particular prey species or have a more generalist diet when fewer options are available (Kassinis *et al*. 2024). This may be visible in the spatial differences in productivity in relation to resource availability or diversity. Whereas, competition for these resources may also be intense, leading to fluctuations in their abundance, occupancy and productivity (Fasce *et al*. 2011; Panuccio *et al*. 2019).

Human activities can have significant impacts on animal populations and their habitats on islands (Agudo *et al*. 2010; Fernández-Gil *et al*. 2023). Habitat modification and destruction, development of infrastructure, pollution, and the introduction of invasive species pose major threats to raptor survival and reproduction (Donázar *et al*. 2016; Ecke *et al*. 2017; Singh *et al*. 2021). Development projects, such as urbanization and infrastructure expansion, can fragment habitats and disrupt breeding and foraging areas (Thiollay & Meyburg 1988; Benítez-López *et al*. 2010). Island habitats can often end up often more fragmented and susceptible to disturbance from human activities and invasive species (Sánchez-Zapata *et al*. 2003; Anadón *et al*. 2010). Fragmented habitats can limit the availability of suitable nesting sites, foraging areas, and dispersal corridors for island raptors (Franke *et al*. 2024).

Golden Eagles (*Aquila chrysaetos*) are widely distributed across the Holarctic (Watson 2011). As apex predators, Golden Eagles are opportunistic but also strongly depend on naturally fluctuating abundances of mountain hares (*Lepus timidus*) and different grouse species in the northern boreal forest (Tjernberg 1981; Moss *et al*. 2012). In highly seasonal environments with harsh winters, eagles may use scavenging opportunities especially when prey abundance is low (Singh *et al*. 2021). Golden Eagles also breed on a few islands across the globe. Island populations of Golden Eagles may exhibit specialized adaptations and behaviors in response to the unique ecological conditions of islands, such as reduced flight, dispersal and migration capabilities. For example, Golden Eagles in Scotland inhabit diverse habitats, including moorlands, mountains, and coastal cliffs, where they rely on prey species such as rabbits, hares, and ground-nesting birds (Fielding *et al*. 2024). Habitat quality, characterized by factors such as prey abundance and suitable nesting sites, strongly influences eagle breeding success and population dynamics (Ogden *et al*. 2015; Fielding *et al*. 2020). Historical persecution, primarily driven by conflicts with game management and concerns over predation on livestock, has been a major threat to Golden Eagles in Scotland (Fielding *et al*. 2024). Genetic factors, such as population structure and genetic diversity have played a role in shaping these populations in Scotland (Fielding *et al*. 2024). On Channel Islands, including Santa Cruz Island and Santa Rosa Island, Golden Eagles nested in rocky cliffs and foraged in diverse habitats, but this population has experienced a significant decline over the past century, primarily due to habitat loss, human disturbance, and predation by introduced species such as feral pigs (Roemer *et al*. 2002; Roemer & Collins 2020). Suitable nesting sites with access to prey-rich areas had been crucial for successful breeding in these areas. Introduced feral pigs have significantly threatened the Channel Islands population by disturbing nesting sites, competing for food, and preying on eggs and nestlings (Roemer *et al*. 2002).

From experience on Scotland and Channel Islands and many other places, it is clear that Island populations are especially vulnerable to anthropogenic changes, both directly and indirectly. Gotland is a large island in the Baltic Sea, a world heritage site, with a resident human population of about 60 000 and a seasonally fluctuating population of about 10 000 inhabitants (Petersson *et al*. 2019). Gotland is also known for its iconic Golden Eagle population (Hogstrom & Wiss 1992). This population on Gotland is regarded as the densest isolated population in the world (Tjernberg 1983). This population is especially interesting because the birds are believed to remain resident on the island even though the distance between the mainland and the island is only about 200 km. Their mainland counterparts, on the other hand, may range over vast areas all over Fennoscandia (Singh *et al*. 2021; Singh *et al*. 2024). These features make this population an interesting ecological system to study for population dynamics of highly mobile species when restricted to an island and achieving high densities. The dominant food item for eagles on the island is believed to be wild rabbits (*Oryctolagus cuniculus*), whose population dynamics are driven by a pathogen (Hogstrom & Wiss 1992). This genetically distinct population holds significant conservation value for the region especially in light of ongoing wind farm development on the island (Balotari-Chiebao *et al*. 2016). The space on the island is shared with an increasing population of the sympatric White-tailed Eagle (*Haliaeetus albicilla*), and a human population dependent on farming, livestock, tourism, forestry and more recently, a steep increase in military activity in the region (Jansson & Dahlberg 1999).

In this study we investigate the patterns and drivers of breeding dynamics of the Gotland Golden Eagle population and factors affecting them. We assess the role of habitat composition, human use of the landscape, resource availability and inter and intraspecific interactions and their relative importance, in determining the spatial patterns of territorial productivity. Due to large habitat heterogeneity on the island, we expect habitat composition and its consequences on food availability to drive spatial variation in productivity. Alternatively, the human driven land use on the island may be the principal driver of productivity instead of natural factors such as prey availability. A dense distribution of Golden Eagle territories and presence of White-tailed Eagle territories in the neighbourhood may entail that competition for space and food drives patterns of productivity instead of habitat composition.

### Study area

Gotland, nestled in the Baltic Sea off the southeastern coast of Sweden (56.9 – 58.0°N, 18.0 – 19.3°E), is a large picturesque island known for its ecological diversity and natural beauty. The island’s terrain is characterized by a blend of pine and deciduous forests, rolling hills, farmlands, and limestone cliffs, supporting a diverse array of plant and animal species (County Administrative Board of Gotland, 2022). Gotland’s coastline is characterized by pristine beaches, rocky shores, and secluded coves that support a thriving ecosystem. The island’s marine environment also serves as an important breeding ground and migration route for various species of birds, including terns, gulls, and waterfowl (Nilsson & Hermansson 2021). The island also contains freshwater lakes and wetlands, providing essential habitat for amphibians, waterfowl, and migratory birds. Despite its relatively small size, Gotland is home to roe deer *(Capreolus capreolus)*, red fox *(Vulpes vulpes)*, hares *(Lepus europaeus)*, hedgehog (*Erinaceus europaeus*), wild rabbits (*Oryctolagus cuniculus*) and numerous species of birds and insects. The roe deer population on Gotland was estimated to be c. 850 individuals in 2012 but recent hunting statistics report that c.5000 individuals were hunted on Gotland in the season of 2019-2020 (Swedish Association for Wildlife and Hunting Management, 2020). About 6800 wild rabbits were hunted on the island during the same season.

Agriculture is the dominant land use on Gotland and vast stretches of farmland are dedicated to cultivating crops such as cereals, potatoes, and vegetables. Additionally, orchards and vineyards contribute to the island’s agricultural diversity. Gotland’s forests, composed mainly of deciduous trees such as oak, pine, birch, and aspen, are managed for timber production, biodiversity conservation, and recreational purposes. Tourism plays a significant role in Gotland’s economy. Gotland is home to several nature reserves, protected areas, and national parks that serve both conservation and recreational purposes. These areas included coastal dunes, freshwater lakes, wetlands, and limestone grasslands. Given its island location, maritime activities such as fishing, shipping, and marine transportation are also common. Fishing ports and harbors support commercial fishing operations, recreational boating, and ferry services connecting Gotland to mainland Sweden and neighboring islands.

### Methods and Data

#### Breeding dynamics

The territory mapping for Golden Eagles is conducted by the County Administrative Board (CAB) based on the national monitoring protocol recommended by the Swedish Environmental Protection Agency and coordinated by the Swedish Museum of Natural History. The monitoring method is based on documenting nesting pairs and their young and the results are therefore reported in the form of occupied territories and nests. The monitoring entails minimum four visits to known territories every year (twice between 1 February and 15 June - Spring Check, and twice within 1st June to 15 September - Summer check) to assess occupancy, breeding attempts, eggs produced, number of young produced, and number survived (Åsbrink & Källman 2023). New territories and nests within them when discovered are mapped and added to the annual monitoring. A rough geographic location representing the estimated centroid of the territory is used as geographic coordinate. The White-tailed eagles (WTE) are monitored in a similar manner.

For this study, we first used the monitoring data on territory occupancy and breeding of Golden Eagles collected between 2002-2023, for whole Sweden and compared patterns on Gotland with mainland Sweden. Between 2021-2023, for known Golden Eagle territories on Gotland, we conducted detailed spatial analysis of breeding parameters and also noted through field observations, which among these territories had an overlapping White-tailed Eagle territory in its immediate neighbourhood (within a 6 km buffer).

#### Habitat composition

Habitat composition within eagle territories was extracted and estimated as proportions, from the Swedish National Land Cover Data, a spatial dataset with 10m resolution (Naturvårdsverket 2019). The 25 landcover classes occurred in the study area and were aggregated to nine classes. These are: (1) Mixed forest: comprising birch, aspen, alder, beech, oak, elm and ash; (2) Coniferous forest: comprising Scots pine and Norway spruce; (3) Water: lakes, rivers and canals; (4) Artificial surface: urban areas, construction sites, and roads; (5) Wetlands: saturated land including marshes and bogs; (6) Arable land; (7) Clear cuts within and outside wetlands; (8) Other open land with and without vegetation; and (9) Coastal. The class ‘Artificial surface’ provides a good estimate of permanent human use infrastructure, i.e. level of human disturbance on the island.

#### Food availability

We estimated food availability as abundance and density of two common potential prey species - European rabbit and Roe deer. First, we estimated their densities within 38 territories through distance sampling conducted via ground-based line transects (mean length 1.962 km) conducted during dusk with a flashlight (handheld 550 Lumen LED-searchlight of model Nightsearcher Trio 550). Surveys were carried out in late October 2023 when the detectability is not obstructed by crops and summer vegetation (Barnes *et al*. 1983), and the species populations are relatively stable (Tittensor 1981). Rabbit populations can be highly aggregated based on local habitat conditions, while roe deer are found along forest edges and in the agricultural fields and pastures. Locations of transects were selected through stratified random sampling, using habitat classes derived from habitat types as strata. The habitat types were based on the NMD dataset used for habitat composition estimation above.

Territories were divided into four strata determined by dominating habitat types (arable land, coastal, coniferous forest, mixed forest or open land). For each stratum, five territories were randomly selected for survey. For selected territory, between two or three suitable candidate transect routes were identified using satellite maps and the NMD dataset in QGIS. Open areas suitable for spotlight surveys on Gotland mainly comprise privately owned farmland not suitable for traversing. Fortunately, public roads and paths are abundant on Gotland. Therefore, candidate transects were allocated along seemingly accessible roads and paths in open habitat types with high detectability like dry and rocky areas with low or no vegetation, arable land, and clearcuts. Straight transect segments were prioritized to avoid double counting. Human settlements were avoided whenever possible. The transects were approximately 2 km long and 200 m wide, each covering 0,4 km^2^. One candidate transect was chosen for survey in each territory. The selection among candidate transects was carried out based on suitability and availability.

Each observation was described by species, number of individuals, time of observation, estimated perpendicular distance from transect, angle from observer, and the location of the observer. As distance measures by the provided rangefinder proved unreliable with fast moving animals, especially during nighttime, perpendicular distance from transect was instead estimated by the observer. All participating observers synchronized their distances estimates and survey techniques before performing surveys. Angle from the observer to the observed subjects was derived using a compass and protractor.

#### Intra and interspecific interactions

To assess the role of intraspecific density dependence on breeding patterns, we estimated the variable - local density i.e. the number of neighboring territories within 10 km around each focal territory and also estimated the reproductive status of four nearest territories each monitored year. For possible impact of interspecific interactions, we included a binary variable (1/0) signifying if a White-tailed Eagle territory overlapped with the Golden Eagle territory as 1 and else 0.

### Data analyses

We first investigated the long-term breeding dynamics for Gotland and mainland for comparison. The population productivity was estimated as the average (± S.E.) of yearly productivity (i.e. number of young fledged/occupied territories/year for the entire population) for the period 2002 - 2023. When a territory was occupied, at least four visits were made to record productivity observing from a distance using binoculars or telescope (x20–85). Productivity was expressed as the number of fledglings per monitored pair, considering those which survived to about 51 days old (Steenhof *et al*. 2017).

#### Spatial and short-term temporal dynamics

In the dataset collected by the CAB Gotland, spatial information was available for most territories along with the number of breeding attempts and nestlings that fledged between 2021 and 2023. Average territory productivity was calculated for each territory with confirmed occupancy for all years between 2021 and 2023. Spatial variation in average productivity among territories (territory productivity) was defined as the total number of young fledged per occupied territory per year between 2021 and 2023. Spatial autocorrelation in productivity was investigated to assess the effect of nearest neighbors on territorial productivity through Moran I and to account for spatial bias (Moran 1950).

#### Drivers of breeding dynamics

We then estimated the values of drivers of productivity dynamics across the island using generalized linear models (with Gaussian distribution), with territory productivity (mean per territory for the three sampling years) as a response variable. The explanatory variables were proportion of different habitat types including the index of human disturbance (through habitat composition with proportion of Artificial surface in the territory), food availability measured as prey density (European hare and rabbit together, and roe deer), and inter and intraspecific interactions, as presence of White-tailed Eagles (overlap of a core of GE territory with a WTE territory – Yes=1/No=0), and local density.

Local density was estimated using ‘dnearneigh’ function in package ‘spdep’ in R, that creates a list and then count of neighbors for each territory within the 10 km radius. For the four nearest neighbours we checked ‘the proportion of them reproducing’ each monitored year and tested with a chi-squared test if the productivity among neighboring territories is dependent on the year (Zar 2010). A significant test result would indicate that reproduction rates differ significantly across years, suggesting that not all neighbors reproduce every year.

Due to a low sample size for which all spatial variables could be extracted, we first fitted univariate models to assess the relationship between each predictor and territorial productivity and then followed with a multivariate model including those predictors which had a statistically significant relationship with productivity. We used AIC based model selection criteria to compare models and used model dredging to identify the best average model (Burnham *et al*. 2011).

## Results

We identified 86 Golden Eagle territories on Gotland, out of which 72% were occupied on an average during the three years of intensive survey. The average population productivity between 2002 and 2023 for Gotland and mainland Sweden was 0.41 (S.E.: ±0.04 and SD:0.37) and 0.46 (S.E.:±0.03 and SD:0.14), respectively (Figure S1). On Gotland it fluctuated between 0.2 and 0.85 while for mainland Sweden, it fluctuated between 0.26 and 0.69. Year 2023 was the most productive year on Gotland, where all territories where adults attempted breeding produced fledglings (Table 1).

**Table 1.** Estimated breeding parameters for Gotland Golden Eagle population between 2021 and 2023.

| Parameter | 2021 | 2022 | 2023 | Mean |
| --- | --- | --- | --- | --- |
| Number of occupied territories on Gotland | 65 | 75 | 77 | 72.33 |
| Number of breeding attempts for population | 28 | 41 | 43 | 37.33 |
| Number of territories not attempting breeding | 37 | 34 | 34 | 35.00 |
| Number of successful breeding attempts | 19 | 32 | 43 | 31.33 |
| Number of failed breeding attempts | 11 | 10 | 0 | 7.00 |
| Number of nestlings on Gotland | 19 | 40 | 58 | 39.00 |
| Breeding attempts per territory | 0.43 | 0.55 | 0.56 | 0.51 |
| Number of breeding attempts | 0.57 | 0.45 | 0.44 | 0.49 |
| Successful breeding per territory | 0.29 | 0.43 | 0.56 | 0.43 |
| Successful breeding per attempt | 0.68 | 0.78 | 1.00 | 0.82 |
| Failed breeding per territory | 0.17 | 0.13 | 0.00 | 0.10 |
| Failed breeding per attempt | 0.39 | 0.24 | 0.00 | 0.21 |
| Fledglings per occupied territory (population productivity) | 0.29 | 0.53 | 0.75 | 0.52 |

### Spatial and short-term temporal dynamics

On Gotland, the number of breeding attempts were variable, ranging between 28% to 43%, out of which 32% succeeded on average (Table 1). The total number of fledglings was highly variable between the three years, ranging between 19, 32 and 43 during 2021, 2022 and 2023 respectively across all the territories. The average productivity on the island was 0.53.

Spatial variability in territory productivity was analyzed for 71 territories which varied between 0 and 2, with an average of 0.87 fledglings per breeding pair, and a CV of 0.6 (Figure 1, Table 1). The territories were evenly distributed across the study area, except the area surrounding the city of Visby, where a noticeable gap exists. Furthermore, the average productivity of three of the four closest territories to Visby ranged between 0 and 0.5. The highly productive territories (productivity = 1.5 – 2) were evenly dispersed across the island indicating a lack of geographic bias in productivity and were also the ones with less variation in productivity (Figure 1). There was no significant spatial autocorrelation observed in the territorial productivity (Moran I= −0.03, p=0.32). The average distance to the nearest neighboring territory on Gotland was about 5.3 km, with a range between 3.70 to 9.60 km.

**Figure 1.**
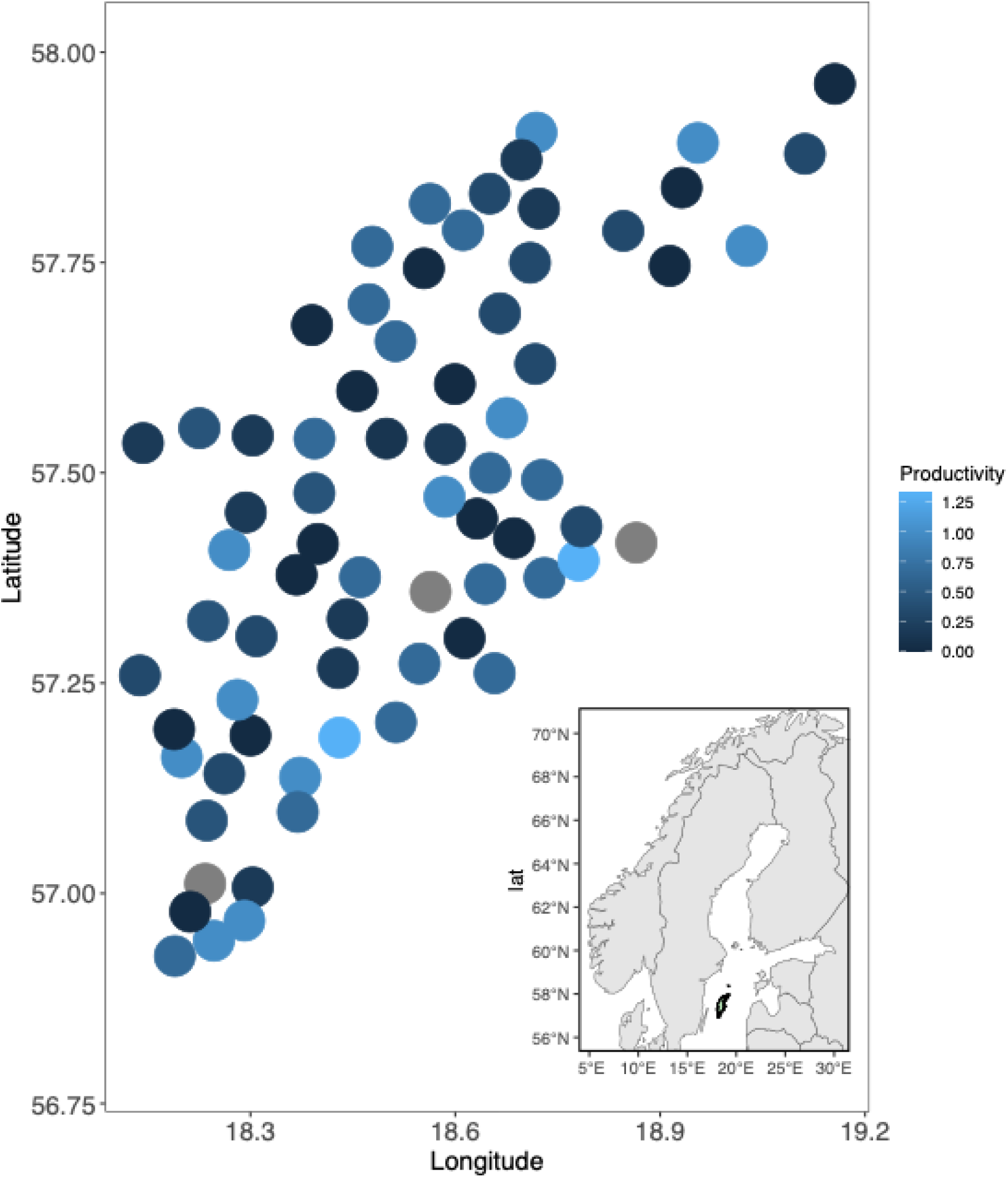
A representation of Golden Eagle territories (n=71, black dots) along with their estimated average productivity (0-2, represented by the colour of the dot) on Gotland during 2021-2023 (the circles are generated around a random point within the territory).

### Correlates of breeding dynamics

Due to a limited sample size (n=86), it was not possible to fit complex multiple variate models due to overfitting and overparameterization, we therefore assessed univariate relationships between productivity and the correlates. Following this, we formulated the additive models using variables that were significant in univariate relationships.

For univariate model with habitat composition as the predictor of productivity, coniferous forest and arable land were the most common habitat type in all territories to varying degrees (Figure 2, 3 & S2). Other open land and mixed forest rarely covered more than 30 % of territory surfaces. Clearcuts, wetland, and inland water existed only in small patches within the territories. The proportion of marine habitat depended on the nearness of the territory to the coast which had a significant amount of conifer forest and arable land (Table 2 & 3, Figure 2, & S1). Therefore, it was positively associated with territory productivity (Table 2 & 3).

**Figure 2.**
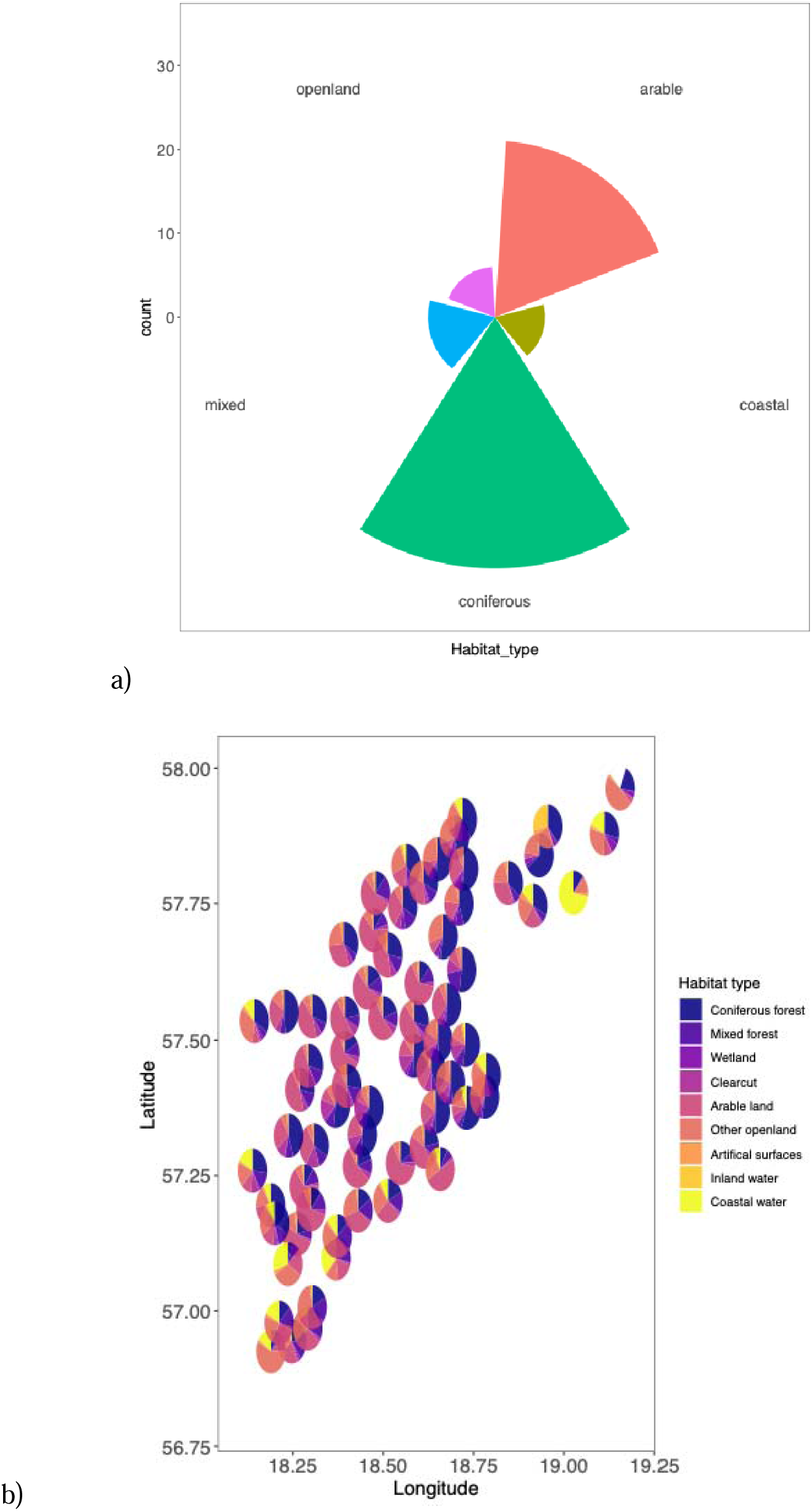
a) Proportion of dominant habitat types in 71 Golden Eagle territories monitored on the Island of Gotland, Sweden between 2021-2023. b) Habitat composition for each territory.

**Figure 3.**
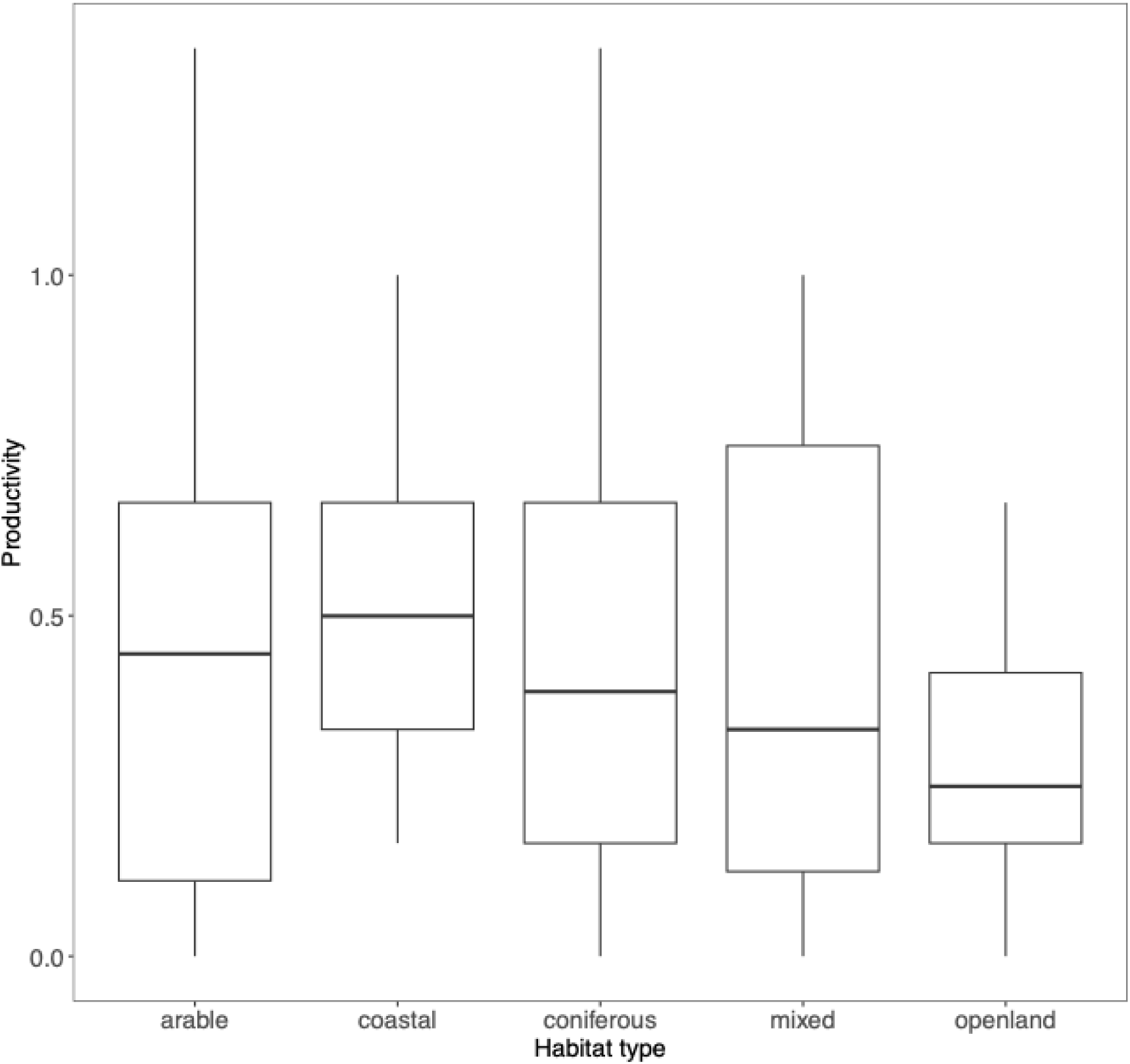
Average productivity of Golden Eagle territories (n=71) dominated by different habitat types (when a single category covered >40% of the surface area) on Gotland, Sweden between 2021-2023. Horizontal bar is the median and the whiskers represent the lower 1^st^ and the upper 3^rd^ quantiles respectively.

**Table 2.**
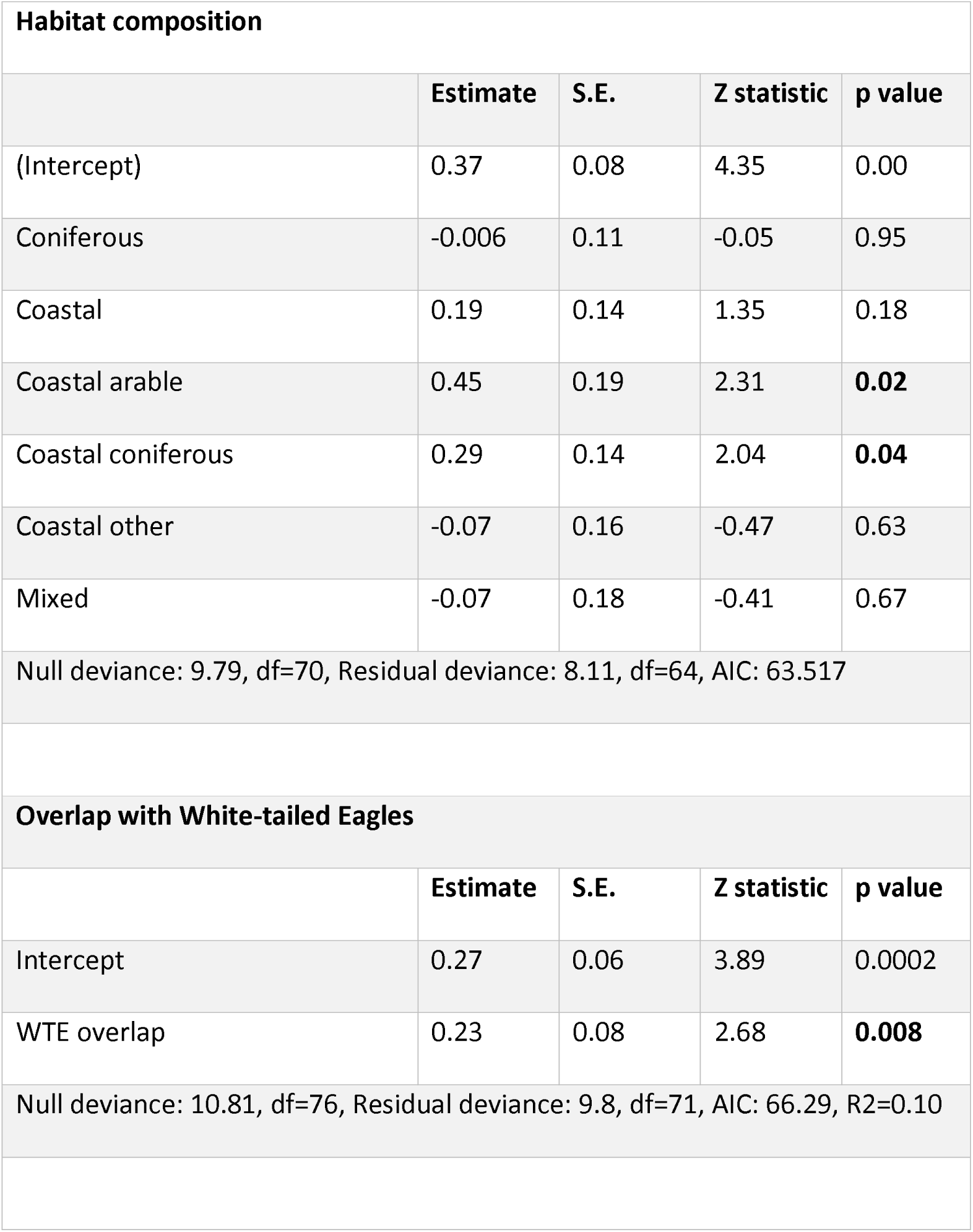
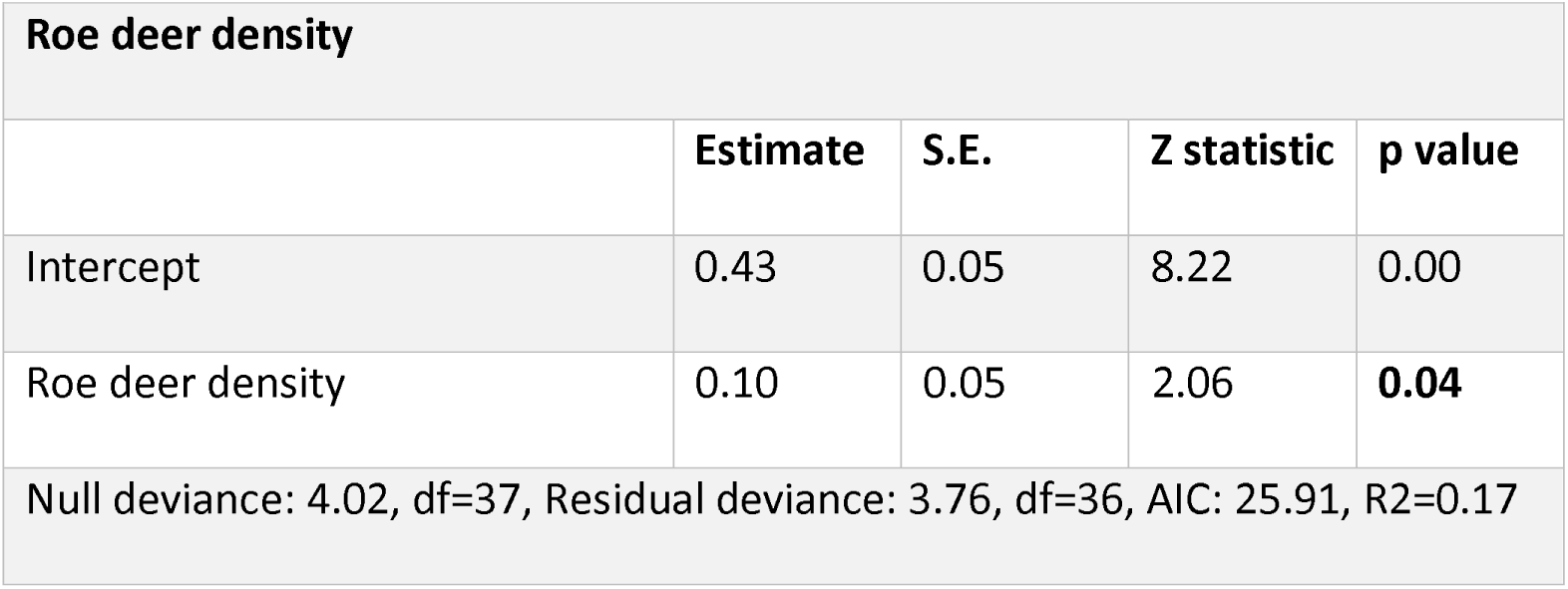
Univariate generalized linear models predicting relationship between territorial productivity and correlates (variables: habitat composition, overlap with White-tailed Eagle territories, Roe deer density) for 86 Golden Eagle territories. Artificial surface is the reference habitat (Intercept) for the habitat categories. All variable values were not always available for all territories.

| **Habitat composition** |  |  |  |  |
| --- | --- | --- | --- | --- |
|  | **Estimate** | **S.E.** | **Z statistic** | **p value** |
| (Intercept) | 0.37 | 0.08 | 4.35 | 0.00 |
| Coniferous | -0.006 | 0.11 | -0.05 | 0.95 |
| Coastal | 0.19 | 0.14 | 1.35 | 0.18 |
| Coastal arable | 0.45 | 0.19 | 2.31 | **0.02** |
| Coastal coniferous | 0.29 | 0.14 | 2.04 | **0.04** |
| Coastal other | -0.07 | 0.16 | -0.47 | 0.63 |
| Mixed | -0.07 | 0.18 | -0.41 | 0.67 |
| Null deviance: 9.79, df=70, Residual deviance: 8.11, df=64, AIC: 63.517 |  |  |  |  |
| **Overlap with White-tailed Eagles** |  |  |  |  |
|  | **Estimate** | **S.E.** | **Z statistic** | **p value** |
| Intercept | 0.27 | 0.06 | 3.89 | 0.0002 |
| WTE overlap | 0.23 | 0.08 | 2.68 | **0.008** |
| Null deviance: 10.81, df=76, Residual deviance: 9.8, df=71, AIC: 66.29, R2=0.10 |  |  |  |  |

| Roe deer density |  |  |  |  |
| --- | --- | --- | --- | --- |
|  | Estimate | S.E. | Z statistic | p value |
| Intercept | 0.43 | 0.05 | 8.22 | 0.00 |
| Roe deer density | 0.10 | 0.05 | 2.06 | **0.04** |
| Null deviance: 4.02, df=37, Residual deviance: 3.76, df=36, AIC: 25.91, R2=0.17 |  |  |  |  |

About 50 WTE territories are known and monitored on the island. The WTE overlap was noted in 43 of the 71 analyzed Golden Eagle territories during the study period (Figure 4). The presence of WTE in Golden Eagle territories was observed across the entire east coast, and the northern and southern edges of the island (Figure 4), but less so in the central and western areas of Gotland. All Golden Eagle territories with high productivity (1.5 – 2) but one, overlapped with WTE presence. The WTE presence positively associated with the Golden Eagle territory productivity (Table 2).

**Figure 4.**
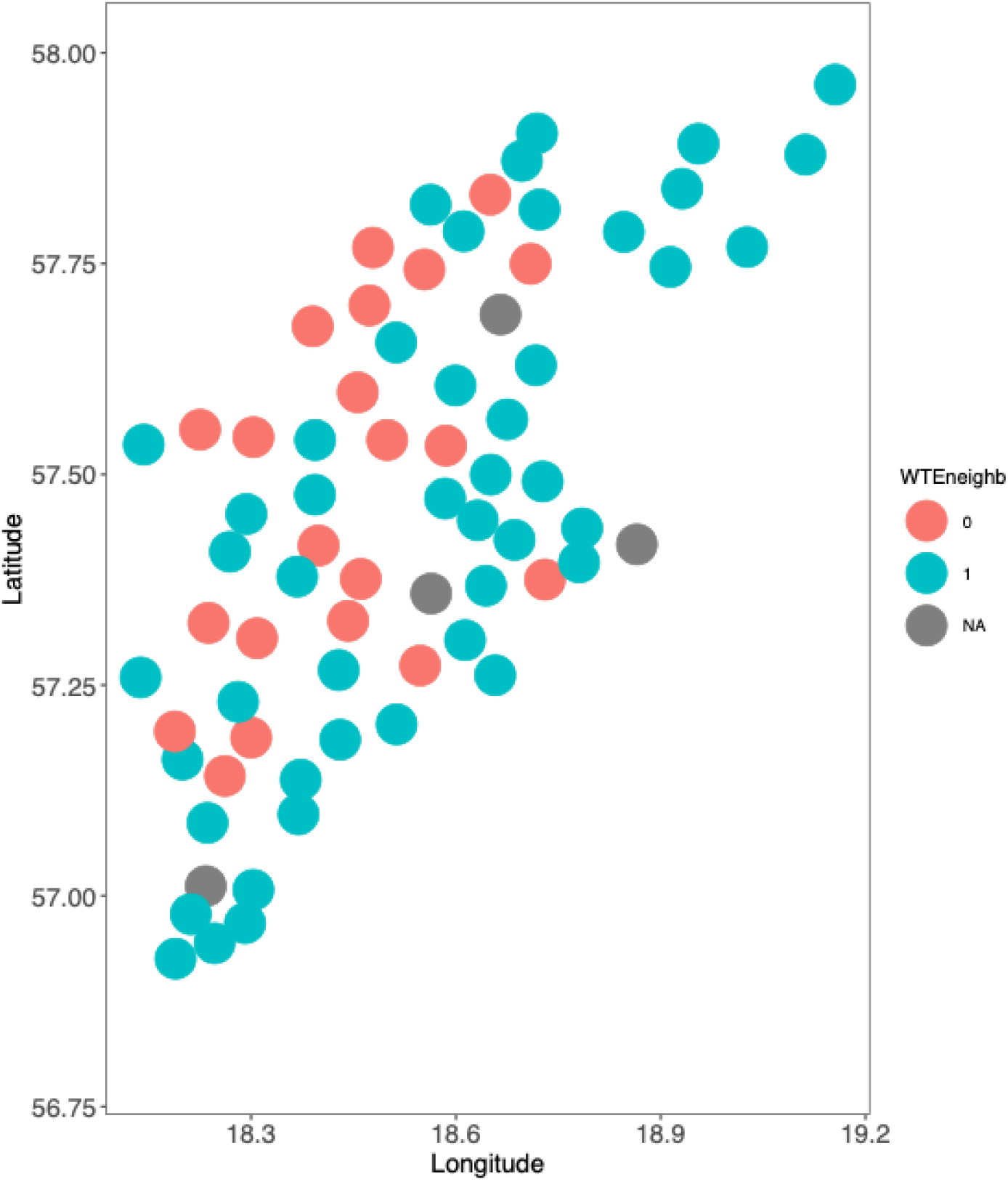
Overlapping Golden Eagle territories ‘1’ with White-tailed Eagle territories on Gotland, Sweden (n=71). The grey territories had no data available.

Roe deer density ranged between 0-0.05/km^2^ within the territories, and (t= 2.06, p<0.05, Table 2 & 3, Figure 5) in the territory had a positive influence on the productivity. The rabbit density was highly variable ranging between 0-0.36/km^2^ at places where aggregations were observed (Figure 5). However, there was no significant effect of rabbit and hare density on productivity.

**Figure 5.**
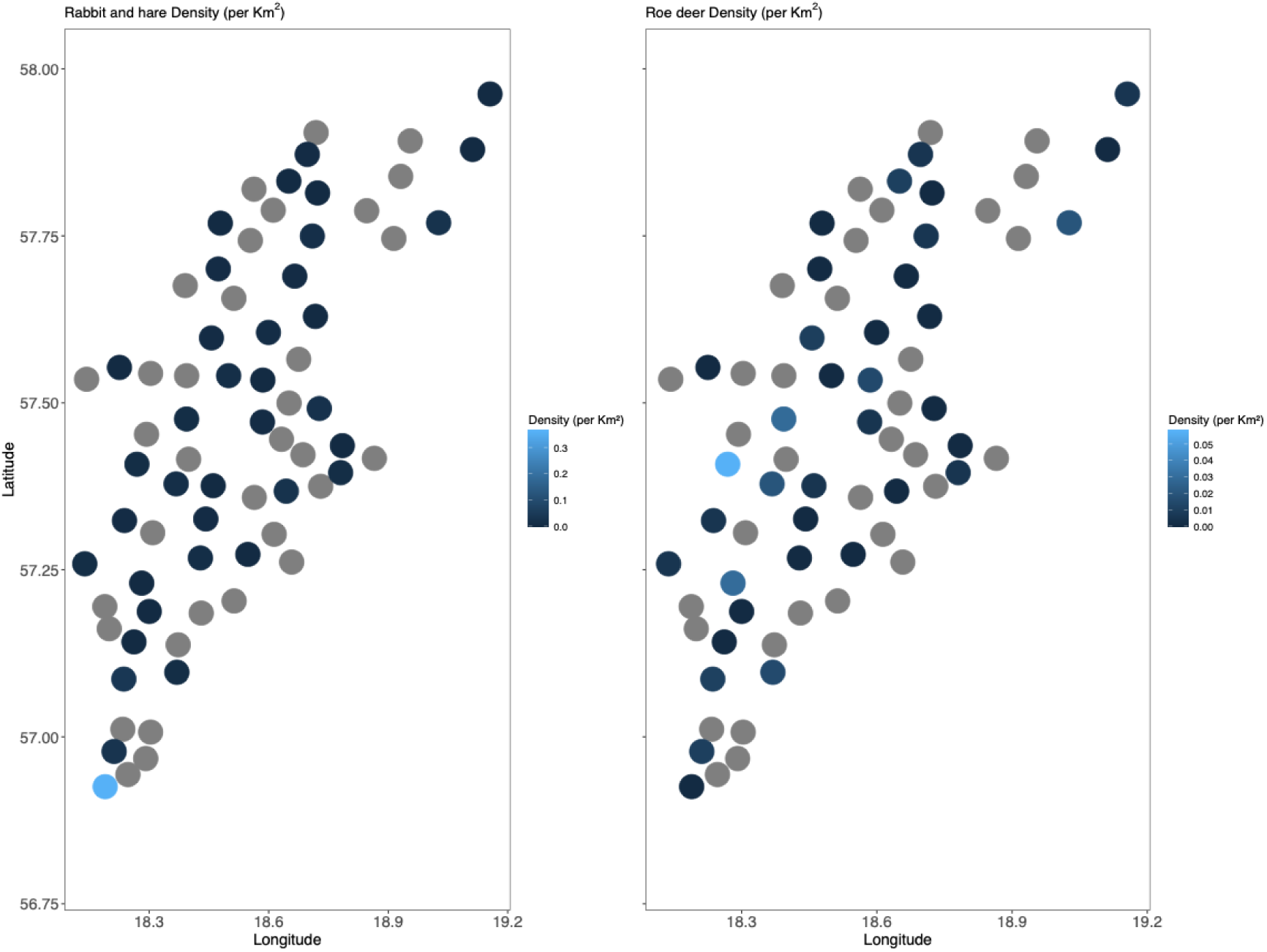
Density (individuals/Km^2^) of a) Rabbits and hares and b) Roe deer in 38 surveyed Golden Eagle territories on Gotland, Sweden, estimated through distance sampling based on line transect method during autumn 2023. Territories in colour are those that were surveyed and the grey ones were not surveyed.

According to multivariate additive models including the above three variables, the roe deer density and coniferous forest and coastal were positively associated with a higher productivity and were retained in the final model following model selection (Table 3). Overlap with the WTE was dropped in the final model.

**Table 3.** Generalized linear models predicting relationship between territorial productivity and correlates (variables: overlap with White-tailed Eagle territories, habitat category, density of prey) for 86 Golden Eagle territories. Results for full additive model (a) and final model (b) are presented. Artificial surface is the reference habitat (Intercept) for the habitat categories.

|  | Estimate | S.E. | Z statistic | p value |
| --- | --- | --- | --- | --- |
| (Intercept) | 0.36 | 0.11 | 3.25 | 0.002 |
| WTE Overlap | -0.02 | 0.14 | -0.18 | 0.85 |
| Coniferous | 0.07 | 0.14 | 0.51 | 0.61 |
| Coastal | 0.09 | 0.19 | 0.49 | 0.62 |
| Coastal Arable | 0.05 | 0.35 | 0.14 | 0.88 |
| Coastal Coniferous | 0.50 | 0.24 | 2.06 | 0.04 |
| Coastal other | 0.06 | 0.22 | 0.28 | 0.77 |
| Roe Deer density | 0.13 | 0.06 | 2.18 | 0.03 |
| AIC=2.43, Residual deviance: 0.26, df = 6, R<sup>2</sup>= 0.75 |  |  |  |  |

b) Final Model
|  | Estimate | S.E. | Z statistic | p value |
| --- | --- | --- | --- | --- |
| (Intercept) | 0.22 | 0.1 | 2.08 | 0.07 |
| Coniferous | 0.23 | 0.15 | 1.57 | 0.15 |
| Coastal | 0.44 | 0.18 | 2.43 | 0.04 |
| Roe Deer density | 0.27 | 0.07 | 3.92 | 0.005 |

AIC=1.46, Residual deviance: 0.29 on df=7, $R^2 = 0.73$

The mean distance between nearest neighboring Golden Eagle territories on Gotland was c. 5380 m (SD = 1239), ranging from 3760 to 9610 m. The two northern most territories on the small island of Fårö were isolated from other territories and were therefore excluded in these analyses (Figure 1). Territorial productivity did not correlate significantly with mean distance to the nearest neighbor and there was no significant effect of local density on the productivity. However, neighbourhood productivity varied significantly across years, suggesting that only a proportion of neighbors reproduced every year (χ2-squared = 127.17, df = 6, p-value < 0.01, Figure 6 & S3).

**Figure 6.**
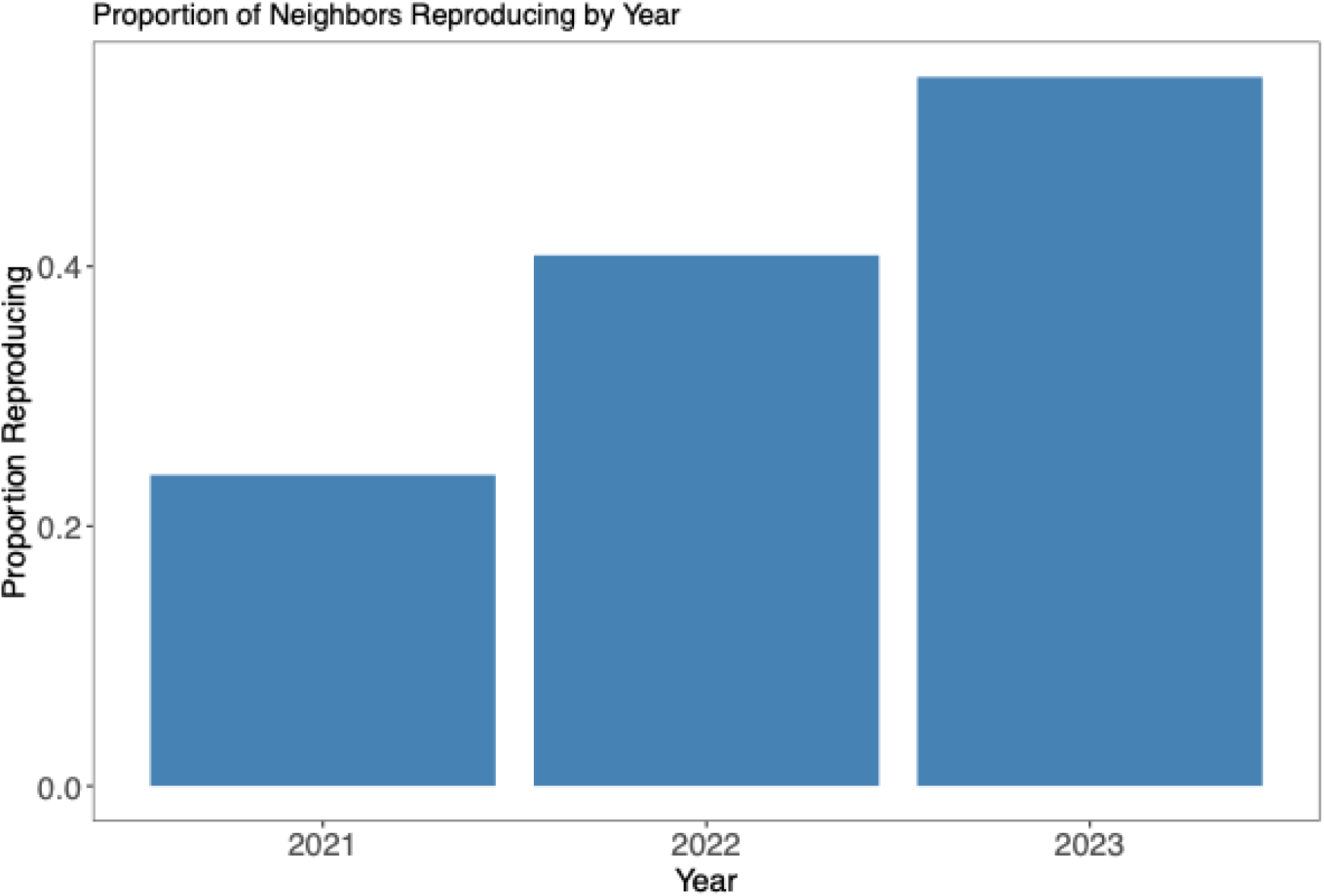
The proportion of neighboring territories (out of n=4) reproducing every year (2021-2023) to each checked Golden Eagle territory (n=71) in Gotland, Sweden, during the three years of monitoring.

## Discussion

We present the first detailed spatial and temporal picture of Golden Eagle territorial dynamics on the island of Gotland, an ecologically and culturally valuable raptor population under pressure from human use and modification of landscape. We reveal several new and interesting results on the ecology of this population, its breeding variability and the factors affecting these.

Compared to mainland Sweden, the population appears to have a similar productivity albeit with a higher interannual variability. This was lower in relation to previous estimates, where Tjernberg (1983) estimated it to be 0.64 for northern Sweden between 1975 and 1980 and also low in comparison to the neighboring Norway (0.58 between 1970 and 1990) (Gjershaug 1996) and Finland (0.64 between 1970 and 1989) (Virolainen & Rassi 1990). In Västerbotten, the northern Swedish county that is the stronghold of the population, the productivity was 0.58 (Trulsson 2019). Despite a low population productivity, the number of territories occupied has increased on Gotland (n=86), potentially a reflection of monitoring effort. On the other hand, it is believed that Gotland eagles don’t leave the island (although not proven), which could indicate, as the population developed, younger birds dispersed and established new territories as the population developed. This has led to an actual increase in the number of territories and hence the population.

When comparing with other main populations across the globe, the north American population’s productivity ranges from 0.7 to 1.1 fledglings per breeding pair annually (Mcintyre & Schmidt 2012; Morneau *et al*. 2012). This varies with habitat quality, particularly in areas like Alaska and western U.S., where some declines have been noted due to habitat loss and prey availability (Millsap *et al*. 2022). Productivity in the Alps, including Italy, varies between 0.6 and 0.8 fledglings per pair annually, with fluctuations linked to environmental conditions such as prey availability and human disturbance (Di Vittori & López-López 2014). In Spain, Golden Eagle populations show relatively high productivity, averaging around 0.9 fledglings per pair per year in well-preserved habitats (Fernández-Gil *et al*. 2023). However, urbanization and agricultural expansion have caused some regional declines. In other parts of Scandinavia, particularly in mainland Sweden and Norway, productivity is moderate, with about 0.7 to 1.0 fledglings per pair annually, depending on prey availability and levels of human disturbance (Gjershaug *et al*. 2018).

Across space, the territories do not follow any special aggregations and are wide spread across the island (Figure 1). This is probably a consequence of the distribution of preferred habitat for nesting (coniferous) and hunting (arable land) (Figure 2). Eagles on Gotland primarily nest in coniferous trees (with some exceptions) and are observed hunting in open habitats. Agriculture is a dominant land use along with forestry on Gotland. Therefore, coniferous forest along with arable land are key limiting (breeding) habitats for the Golden Eagles there (Sandgren *et al*. 2014; Singh *et al*. 2016). These habitat classes were evidently significant in our habitat composition analyses both in the univariate and multivariate models. Eagle are opportunistic feeders and may adapt their diet to the available food which the island of Gotland has in plenty through the diversity of birds, rabbits and hare, roe deer, fish and other mammals (Tjernberg 1981; Hogstrom & Wiss 1992; Clouet *et al*. 2017). Territories close to coast that also had a high proportion of coniferous forest and arable land had higher productivity (Figure 2). Occupying territories closer to the coast likely provides access to both land based and marine prey (primarily breeding waterfowl) and hence positively influences productivity, which is contrary to what has been reported in the island of Sardinia (Di Vittorio *et al*. 2020). These differences are probably due to a much steeper terrain and high human population density in Sardinia. Similar observations were found for Sicily (Di Vittori & López-López 2014).

Another interesting aspect that emerged was the positive association between Golden Eagle productivity and territorial overlap with White-tailed Eagle territories. The White-tailed Eagles are generally known to rely on aquatic fauna more than Golden Eagles (Sulkava *et al*. 1997; Ekblad *et al*. 2020), which would entail that White-tailed eagle territories are likely to be more abundant towards the coastal areas. This was clearly the case as the overlap between the two species was often consistent towards the coast (Figure 4) and yet also often occurred throughout the island. Both the Golden and the White-tailed Eagles are generalists and scavengers, and can range over vast areas (Ekblad *et al*. 2016; Balotari-Chiebao *et al*. 2018; Singh *et al*. 2021). Hence it is not a surprise that the relatively small island of Gotland is entirely available to both the species and therefore, the positive effect of the overlap with the White-tailed Eagle was more an artefact of greater feeding opportunities for Golden Eagles towards the coast rather than to do with the overlap with the White-tailed Eagles. This was again validated in the positive effect of coastal habitat on productivity above.

The density of roe deer emerged as a significant predictor of increasing productivity and not the rabbits. This was unexpected as the general belief that has existed in the region and among the ornithologists is that this Gotland Golden Eagle population is sustained through a high density of rabbits (Tjernberg 1983; Hogstrom & Wiss 1992). We found that rabbits were distributed too heterogeneously and occurred in aggregated areas where sand dunes were abundant. This could be a reason that their effect on the overall scale of the island was not evident. Although in some years with high density they may become more dominant. Eagles are known to prey on roe deer elsewhere but have not been studied or reported in Sweden (Pedrini & Sergio 2001; Sidiropoulos *et al*. 2022). Roe deer give birth precisely during the breeding season of Golden Eagles, i.e. from late April to early July (Jarnemo *et al*. 2004) and roe deer fawns therefore become potentially an important prey for eagles during this critical period. Simultaneously we also discovered based on the roe deer hunting statistics on Gotland, that c.5000 individuals were hunted on Gotland just within the season of 2019-2020 (Figure S4, Swedish Association for Wildlife and Hunting Management, 2020), while c.6800 wild rabbits were also hunted on the island during the same season. For roe deer these figures present a stark reality that their population has rapidly increased on Gotland just within 10 years which potentially also explains the general rise in the number of nesting Golden Eagle pairs during the last 10 years. Clearly, the role of roe deer in affecting eagle population dynamics, needs further investigation and our study lays a path for that.

Another interesting aspect that emerges from this study is the role of neighbours. We began with exploring signs of potential density dependence on breeding through distances between nearest neighbours or local territory density, as has been tested elsewhere (Fasce *et al*. 2011; Chambert *et al*. 2020; Millsap *et al*. 2022). We did not find a negative effect of a higher number of neighbours or distance to the nearest neighbours on productivity, but we did find that the proportion of neighbouring breeding territories varied between years, indicating some influence of local density on breeding (Figure 6). Also, this might be in turn regulated through the weather conditions in the breeding year which are known to significantly affect eagle reproduction (Fasce *et al*. 2011; Fernández-Gil *et al*. 2023; Fielding *et al*. 2024). Our dataset was too small to explore the trends in productivity yet. But the spatial fluctuations in breeding status of territories and their neighbours may allow a dense population to prevail by switching breeding across years.

Intraspecific competition has been regarded as the major driving force in determining the spacing of territorial Golden Eagle pairs in South eastern Spain (Sánchez-Zapata *et al*. 2000). The average distance to the nearest neighboring territory on Gotland was about 5.3 km, with a range between 3.70 to 9.60 km, and the highly productive territories were quite evenly dispersed across the island (Figure 1). The inter-territorial distance between Golden Eagles has been observed to vary by habitat type, availability of nesting sites, prey availability and population density (Pedrini & Sergio 2001; Whitfield *et al*. 2022). In this aspect, Gotland seems to resemble Scotland and Western United States, where distances between neighboring territories have been observed to be between 6 to 10 km at high densities (Whitfield *et al*. 2022). Therefore, in terms of numbers based on average distance, it may seem that Gotland has a high density of territories. However, an important that warrants further investigation is the effect of weather patterns before and during the breeding season which may have influenced the reproductive success, and consequently may explain the year-to-year differences we observed. Further studies need to account for this cause of variation on breeding dynamics (Clouet *et al*. 2017; Franke *et al*. 2024).

Despite of the new knowledge acquired during this study, a number of key questions still remain to be answered about this unique population for its long-term sustainability. Is this population strictly resident on the island and if there is any genetic exchange with the mainland population? (Ferrer *et al*. 2011). A preliminary study with a small sample size showed that the Gotland population was genetically distinct from the mainland population, but the diversity within the population is not known (Näsman 2018). Further, is inbreeding an issue on the island? What is the turn-over rate in and across the territories? Similarly, we lack information about the spatial scales of movements of different age classes, including their annual and seasonal range, dispersal distances and timings, size of territories and home ranges. Such factors are crucial in influencing territorial breeding dynamics, survival, age of maturity and first reproduction (Ferrer *et al*. 2011).

There is also little information on the diet of this population especially in light of our new results and at high densities. Last published studies come from period when the population density was much lower (Tjernberg 1981; Hogstrom & Wiss 1992) and White-tailed Eagle population was at its lowest. Further investigations are needed to quantify the carrying capacity of the island with respect to the available extent of nesting sites and prey species, especially because forestry and hunting are dominant land-uses on the island. Should roe deer populations continue to be hunted at the current levels, more attention must be paid to assess their role in the ecosystem dynamics of the island. There is also immense pressure to develop wind energy on the island, and even though the wind farm distribution is currently limited to the vicinity to a few territories, new construction plans and new permits are regularly proposed. Overall, we highlight many key aspects that need to be urgently addressed to ensure the long-term persistence of this unique population.

Besides the above novel insights emerging from this study, a number of caveats remain which needs attention for future studies. Our analyses were made on a circular scale of 5 km using an assumed centroid of a territory instead of a nest. Territories are often not circular and right shapes deduced using GPS tracking data may allow for better inference of habitat composition and use by the eagles and thereby more realistic analyses. However, it is often not possible to tag a major part of the population and extrapolations will have to be made in that regard. Moreover, territories often merge and split over years and the number of territories is always dynamic depending upon the population density, resources, age structure, turn-over rates and weather conditions driving mortality (Monzón & Friedenberg 2018; Chambert *et al*. 2020). Therefore, conclusions made at a later point may change according to the actual state of the ecosystem. In addition, we lack basic information on the age of the territorial birds, proportion of sub adults and juveniles in the population and survival rates of different age classes, – all of which may significantly impact the findings. Despite these limitations, we have maximized the utility of the limited data available to make basic inferences and lay the path for future studies on the ecology of this population, as well as comparisons with eagles and other raptors elsewhere.

## Acknowledgements

The study was funded through the project Gotlands Örnar by Carl Tryggers Stiftelse, the Swedish Environmental Protection Agency, Naturvårdsverket and the EU - Interreg Aurora Project-Aquila North Grant No. 20366483. We thank the County Administrative Board of Gotland for supporting data collection and research time for AJM.

## Conflict of interest

The authors declare no conflict of interest.

## Author contributions

NS and JM conceptualized the study. JM, RL and AJM collected data and complemented the monitoring. NS and RO conducted the analyses and all authors contributed towards writing of the Manuscript.

## Statement on inclusion

Our study brings together authors from a number of different countries (Sweden, Spain and Netherlands), including scientists and practitioners based in the country (county administrative board of Gotland) where the study was carried out. All authors were engaged early on with the research and study design to ensure that the diverse sets of perspectives they represent was considered from the onset. Whenever relevant, literature published by scientists from the region was cited; efforts were made to consider relevant work published in the local language.

## Data availability statement

The raw data used in the study is confidential due to the protected nature of the species and the population. It can be made available on request and after an appropriate confidentiality agreement is signed with the owners.

## Supplementary material S1

**Figure S1.**
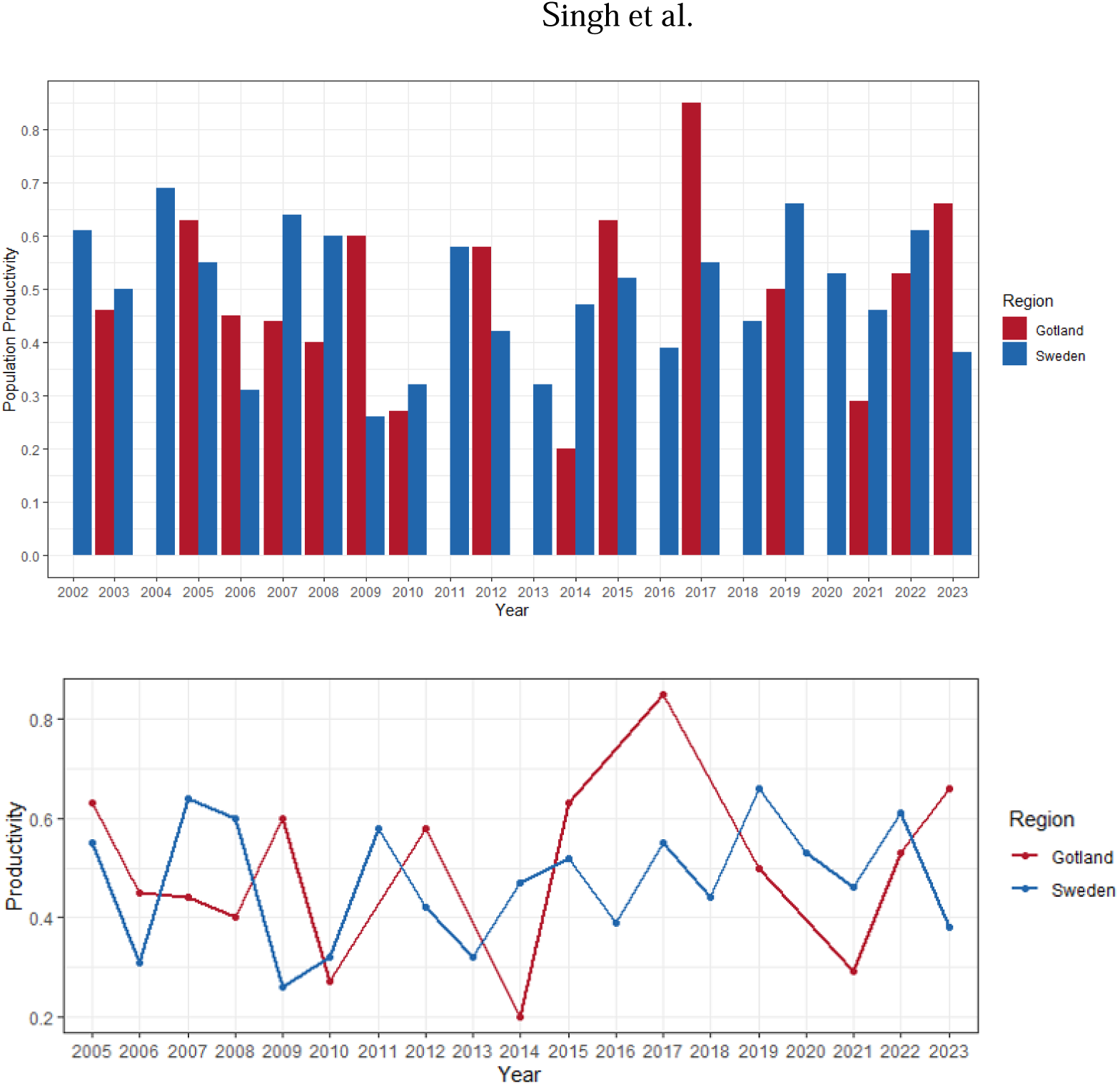
The comparison of population productivity for Gotland and mainland Sweden between 2002 and 2023. Data is missing for Gotland during years 2002, 2004, 2011, 2018, and 2020. Data on breeding dynamics was collected by volunteers and staff of the county administrative boards of Sweden and compiled by the Sweden Museum of Natural History.

**Figure S2.**
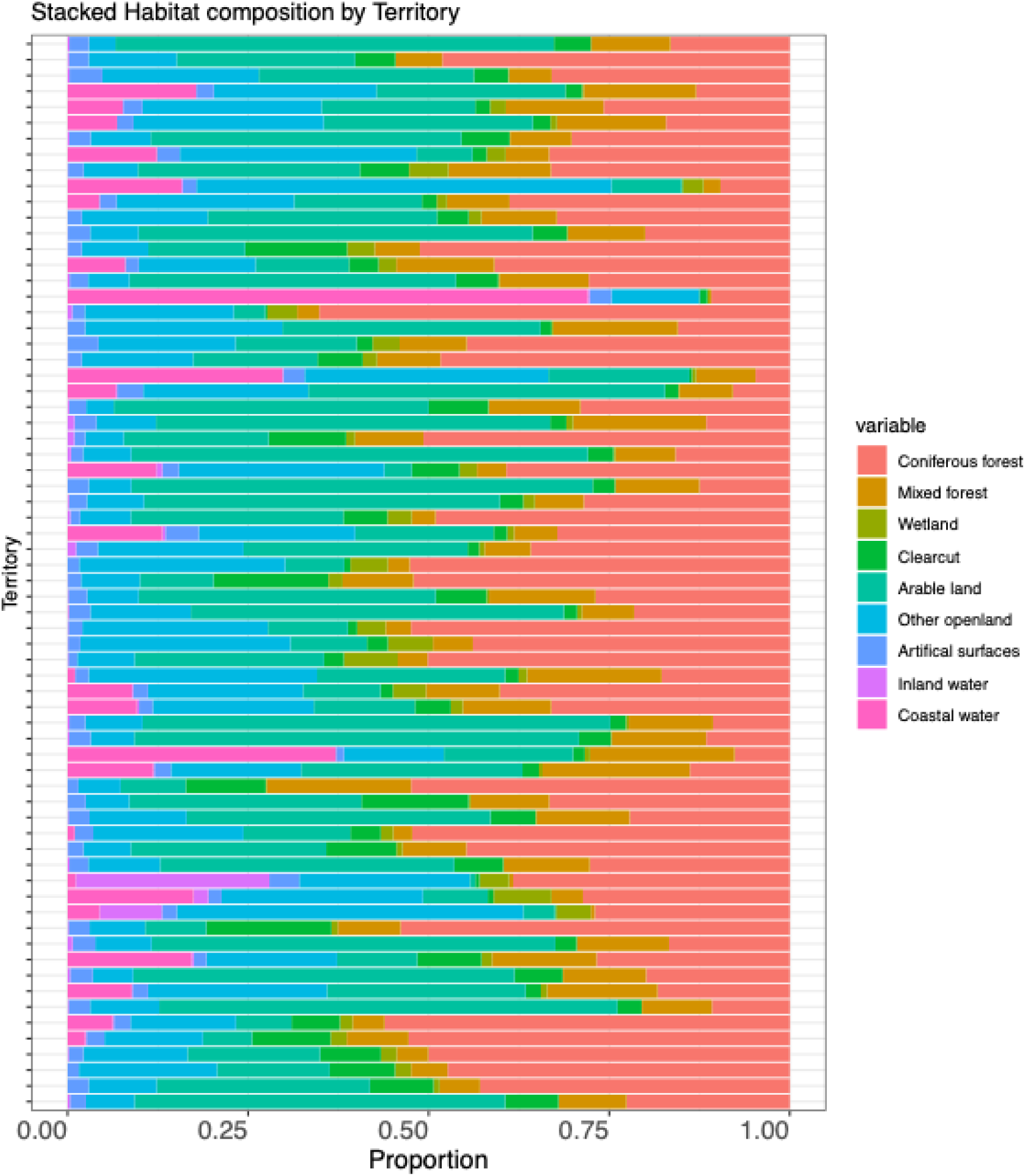
Habitat composition by each monitored Golden Eagle territory on Gotland, Sweden during 2021-2023.

**Figure S3.**
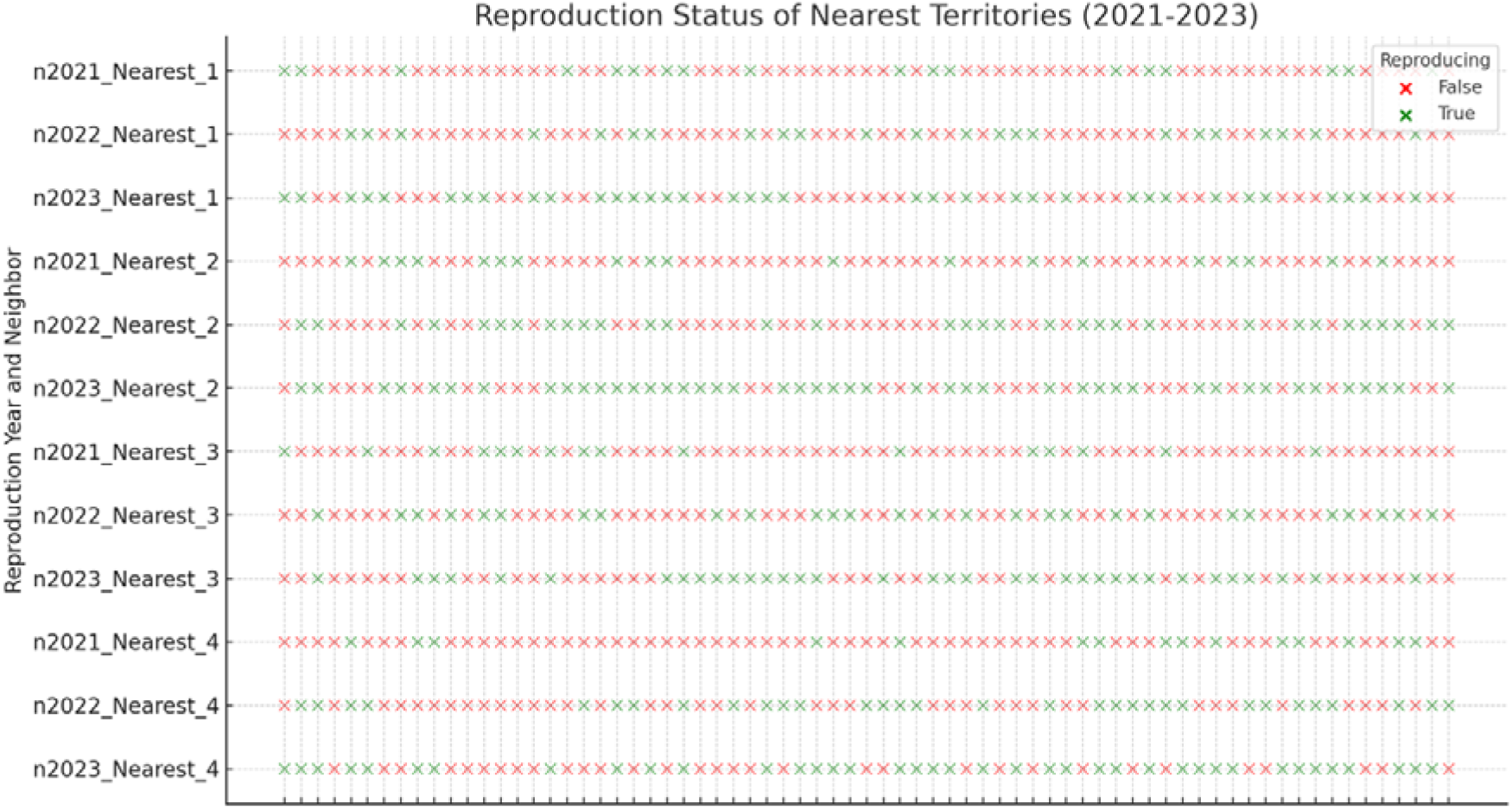
Breeding status of four nearest territories to every monitored Golden Eagle territory on Gotland between 2021-2023.

**Figure S4.**
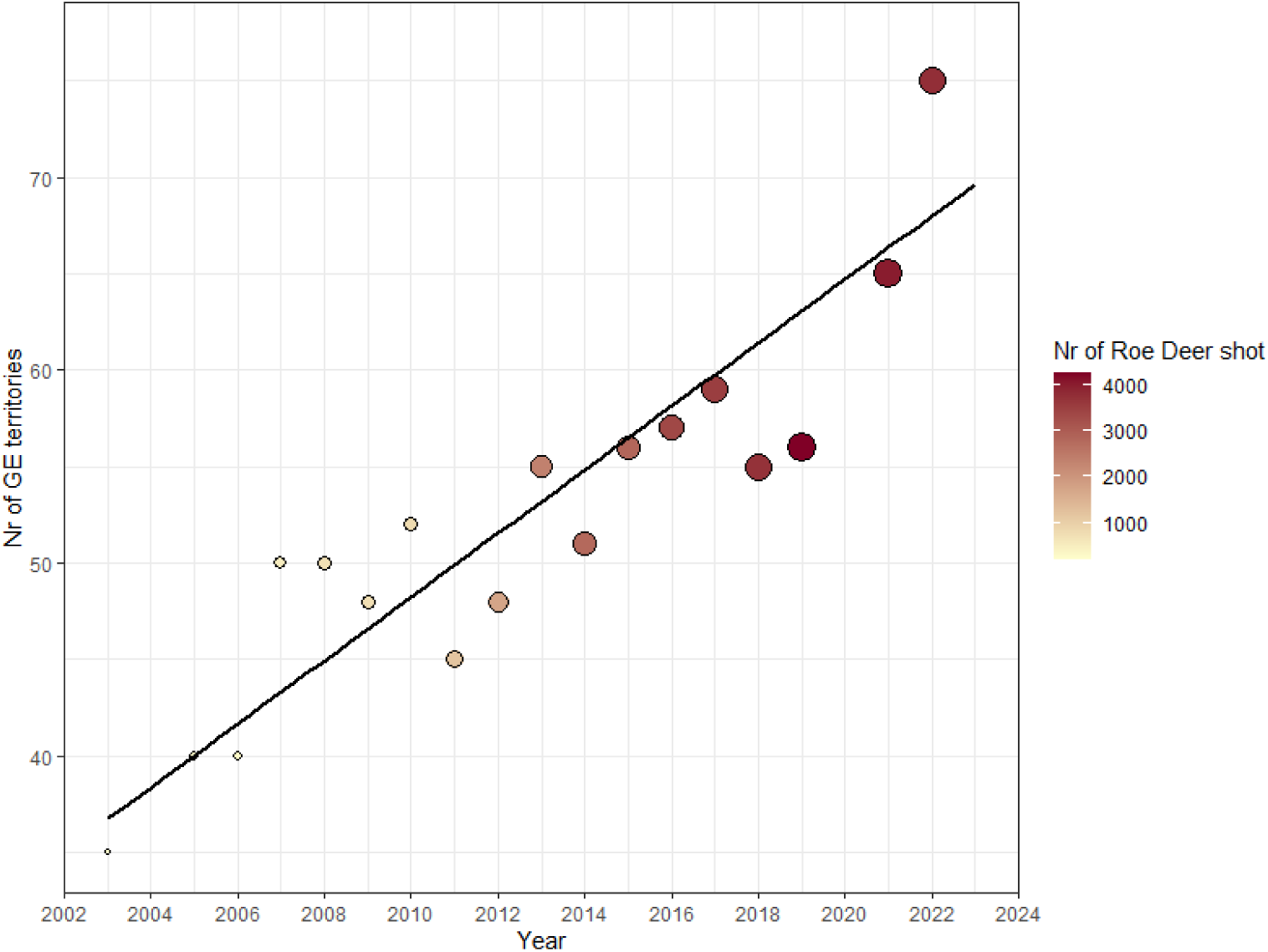
The number of Roe deer shot between 2002 and 2023 on Gotland and the simultaneous development of Golden Eagle population (number of monitored territories).

## Notes

### Competing Interest Statement

The authors have declared no competing interest.

### Summary of Updates

All Figures have been updated to adapt to prevent any issues about confidentiality of data being revealed.

