## Supplementary File for "Island population demography: Breeding dynamics and drivers of Gotland’s iconic Golden Eagles"

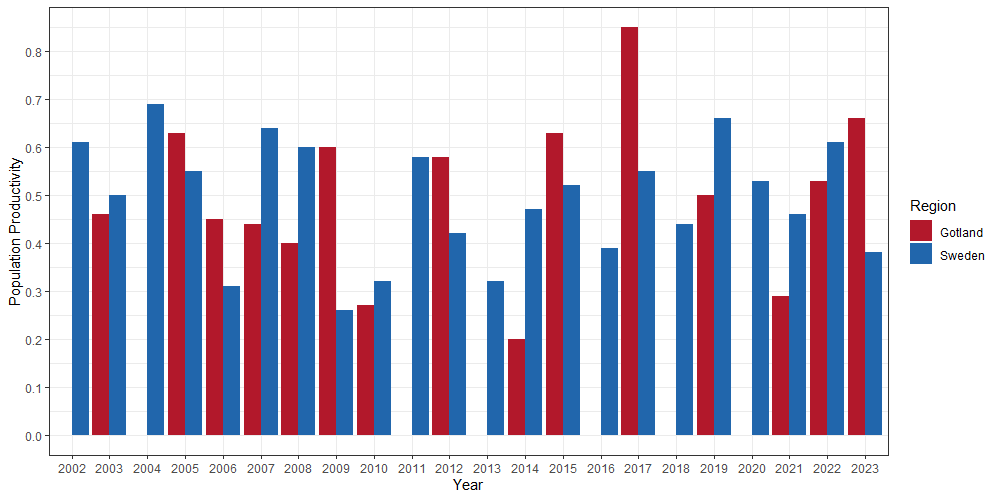

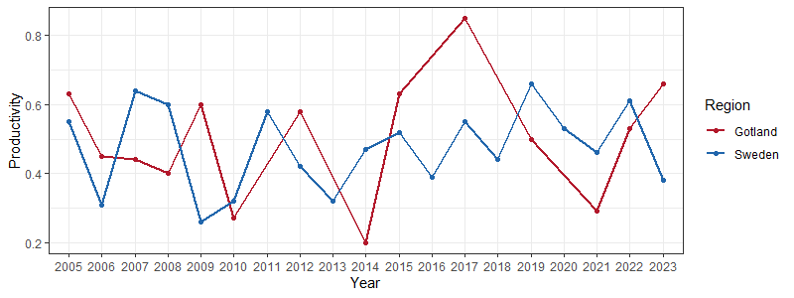

**Figure S1**. The comparison of population productivity for Gotland and mainland Sweden between 2002 and 2023. Data is missing for Gotland during years 2002, 2004, 2011, 2018, and 2020. Data on breeding dynamics was collected by volunteers and staff of the county administrative boards of Sweden and compiled by the Sweden Museum of Natural History.

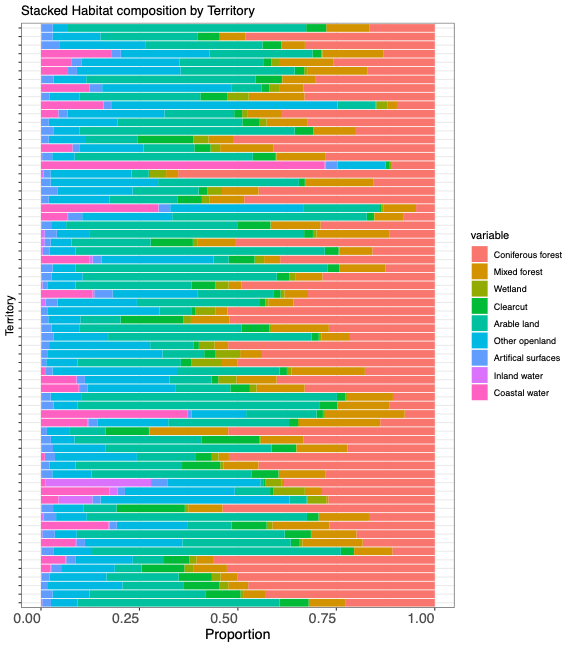

**Figure S2**. Habitat composition by each monitored Golden Eagle territory on Gotland, Sweden during 2021-2023.

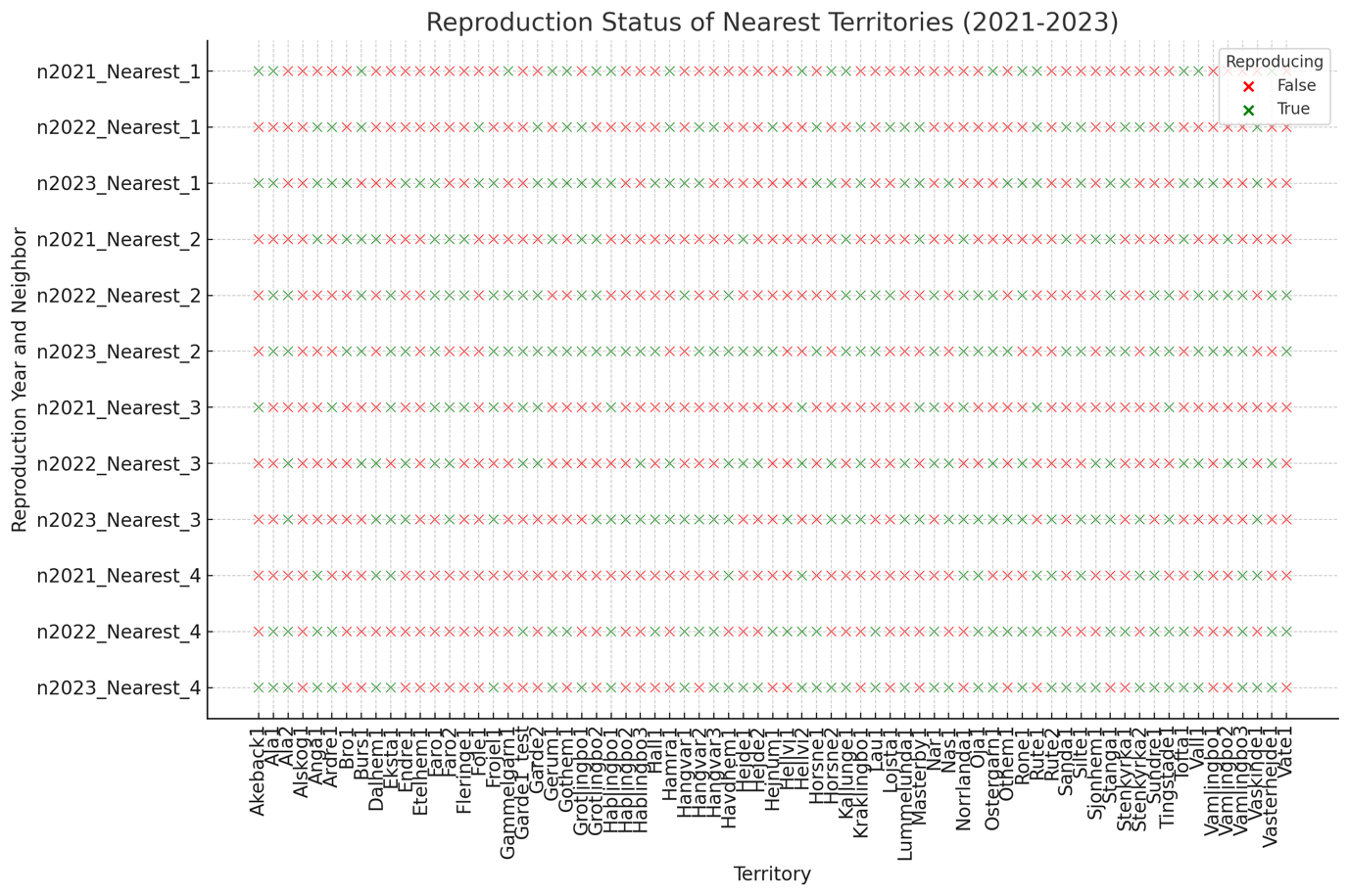

**Figure S****3**. Breeding status of four nearest territories to every monitored Golden Eagle territory on Gotland between 2021-2023.

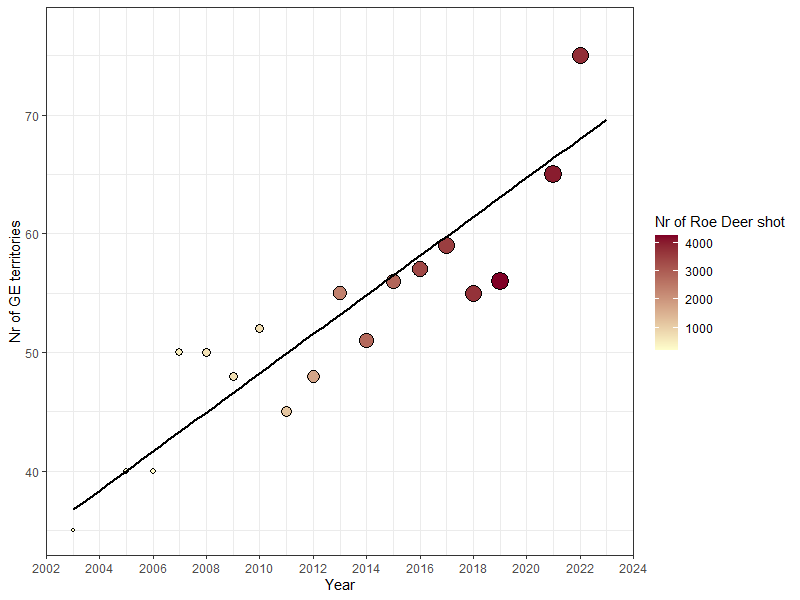

**Figure S4**. The number of Roe deer shot between 2002 and 2023 on Gotland and the simultaneous development of Golden Eagle population (number of monitored territories).
